# Anthrax Toxin Translocation Complex Reveals insight into the Lethal Factor Unfolding and Refolding Mechanism

**DOI:** 10.1101/2020.04.20.049601

**Authors:** Alexandra J Machen, Mark T Fisher, Bret D Freudenthal

## Abstract

Translocation is essential to the anthrax toxin mechanism. Protective antigen (PA), the translocon component of this AB toxin, forms an oligomeric pore with three key clamp sites that aid in the efficient entry of lethal factor (LF) or edema factor (EF), the enzymatic components of the toxin, into the cell. LF and EF translocate through the PA pore (PA_pore_) with the pH gradient between the endosome and the cytosol facilitating rapid translocation *in vivo*. Structural details of the translocation process have remained elusive despite their biological importance. To overcome the technical challenges of studying translocation intermediates, we developed a novel method to immobilize, transition, and stabilize anthrax toxin to mimic important physiological steps in the intoxication process. Here, we report a cryoEM snapshot of PA_pore_ translocating the N-terminal domain of LF (LF_N_). The resulting 3.3 Å structure of the complex shows density of partially unfolded LF_N_ near the canonical PA_pore_ binding site as well as in the α clamp, the Φ clamp, and the charge clamp. We also observe density consistent with an α helix emerging from the 100 Å β barrel channel suggesting LF secondary structural elements begin to refold in the pore channel. We conclude the anthrax toxin β barrel aids in efficient folding of its enzymatic payload prior to channel exit. Our hypothesized refolding mechanism has broader implications for pore length of other protein translocating toxins.

**Significance Statement:** Toxins like the anthrax toxin aid bacteria in establishing an infection, evading the immune system, and proliferating inside a host. The anthrax toxin, a proteinaceous AB toxin secreted by *Bacillus anthracis*, consists of lethal factor and protective antigen. In this work, we explore the molecular details of lethal factor translocation through protective antigen pore necessary for cellular entry. Our cryo electron microscopy results provide evidence of lethal factor secondary structure refolding prior to protective antigen pore exit. Similar to the ribosome exit tunnel, the toxin pore channel likely contributes to native folding of lethal factor. We predict other AB toxins with extended pores also initiate substrate refolding inside the translocon for effective intoxication during bacterial infection, evasion, and proliferation.

## Introduction

The anthrax toxin is not only a deadly *Bacillus anthracis* virulence factor, but also serves as a model system of protein translocation and as a peptide therapeutic delivery platform (1, 2). It’s biological importance and biotechnology utility have spurred significant biochemical and biophysical advances in understanding the anthrax intoxication mechanism. In order to gain entry into the cell, this archetypical AB toxin must cross the endosomal membrane. Membrane penetration is accomplished by the B component of anthrax toxin, termed protective antigen (PA). PA forms a translocon pore through which lethal factor (LF) or edema factor (EF), the A component, translocate. Here, we developed an approach to elucidate the structural and mechanistic details of the anthrax toxin during translocation in an effort to understand how LF unfolds in the endosome, translocates through PA, and refolds in the cytosol.

An overview of the anthrax toxin mechanism has been reviewed by the Collier lab (1) and is briefly summarized here. The first step in intoxication is the 85 kDa monomeric PA binding to host cell receptors. Then the pro-domain of PA is cleaved leaving the 63 kDa PA to oligomerize into heptameric or octameric prepore (PA_prepore_) (3, 4). Up to three LF and/or EF components can bind to the PA_prepore_ heptamer (4–6). The AB toxin complex is endocytosed through clathrin mediated endocytosis (7). As the endosome acidifies, PA_prepore_ undergoes a conformational change to a pore (PA_pore_) (8). This pore inserts into the endosomal membrane to form a channel. The low pH of the endosome and the pH gradient between the endosome and the cytosol facilitate LF or EF to unfold and rapidly translocate into the cytosol in a hypothesized Brownian ratchet mechanism (9). Natively refolded LF and EF in the cytosol are then able to perform their virulent enzymatic functions (10).

The overall structure of the PA_pore_ translocon can be divided into two regions: the funnel and the channel (**Fig. 1A**). The first region, the funnel, facilitates binding and unfolding of LF. LF binds to the rim of the PA_pore_ funnel and is guided down the narrowing structure. The second region of PA_pore_ is the channel, a β barrel that extends from the funnel and spans the endosomal membrane. Three nonspecific PA_pore_ clamp sites (α, Φ, and charge clamp) aid in the translocation of LF (**Fig. 1A**). The α clamp is located at the PA funnel rim, is formed by adjacent PA protomers, and binds helical portions of LF to position them towards the pore lumen. Heptameric PA_pore_ has seven potential α clamp binding sites. A crystal structure of the N-terminal domain of LF (LF_N_) bound to the PA_prepore_ revealed the α clamp binding site, but was unable to resolve the 28 amino acids of LF_N_ passed the α clamp (11). The second clamp site is the Φ clamp, a ring of seven phenylalanine residues that maintain the pH gradient between the endosome and the cytosol (12). The cryoEM structure of apo PA_pore_ revealed the Φ clamp forms a narrow 6 Å diameter ring (13). Secondary structural elements, such as α helices, are too wide to fit through this narrow seal. Therefore, it is hypothesized that peptide substrates must completely unfold and refold in order to translocate through the PA_pore_ and enter the cytosol of the cell (13). The Φ clamp also assists in the unfolding of LF as an unfolding chaperone (2). The third clamp site, the charge clamp, is located within the β barrel of PA_pore_. The charge clamp deprotonates acidic side chains of LF and ensures unidirectional movement of the polypeptide (14, 15). Interestingly, the diameter of the PA_pore_ β barrel is large enough to accomidate an α helix, which would allow for initial refolding to occur inside the pore prior to LF entering the cytosol. However, it remains unclear what structural state LF is in when interacting with the charge clamp and within the β barrel channel.

**Fig. 1.**
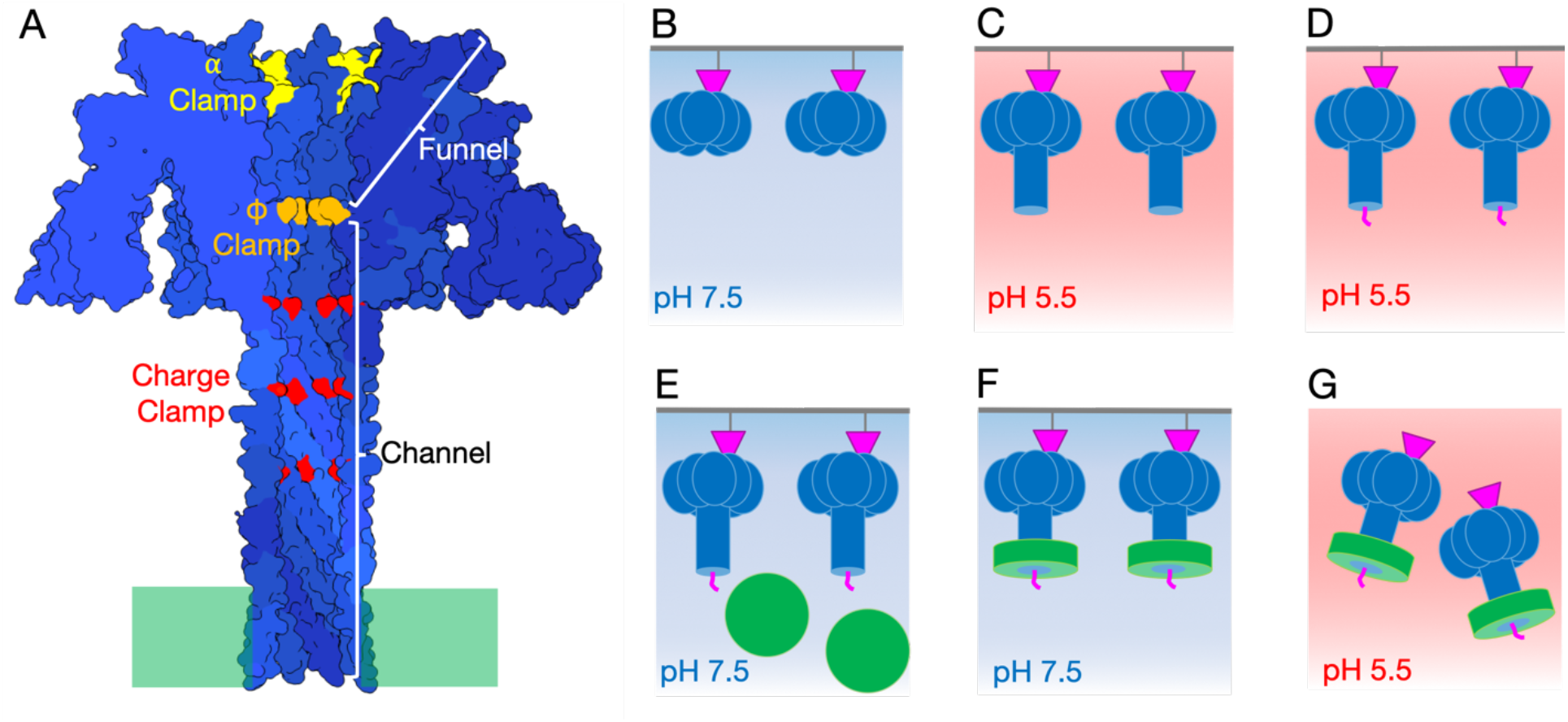
Anthrax toxin immobilization, translocation, and nanodisc stabilization (TITaNS) method. (A) PA_pore_ side view slice with funnel shape from α clamp (yellow) to Φ clamp (orange) and charge clamp (red) inside pore β barrel channel indicated. (B) Immobilization of LF_N_ (magenta) PA_prepore_ (blue) complexes on thiol sepharose beads (grey surface). (C) PA_prepore_ transitioned to PA_pore_. (D) Predicted translocation complex of LF_N_-PA_pore_ at low pH. (E) Addition of pre-nanodisc micelle (green) to complex. (F) Nanodisc formation. (G) LF_N_-PA_pore_-Nanodisc translocation complexes at pH 5.5 on cryoEM grid.

One of the many challenges in studying the anthrax toxin is that it is a dynamic membrane protein that functions under acidic conditions. Thus many questions remain, such as what path LF travels down the endosomal pore lumen from the α clamp to the Φ clamp, whether the Φ clamp adopts multiple states during translocation, and whether LF can partially refold inside the β barrel pore. To address these questions, we developed a novel toxin immobilization, translocation, and nanodisc stabilization (TITaNS) method in combination with cryoEM to structurally characterize PA_pore_ translocating the N-terminal domain of LF (LF_N_). This approach provides unique mechanistic insight into how LF_N_ interacts with the three clamp sites of PA_pore_. We observed density consistent with LF_N_ unfolding prior to the α clamp, translocating through the dynamic Φ clamp, and beginning to refold in the channel of the PA_pore_.

## Results

### Assembly of Anthrax Translocation Complexes

*In vivo*, the anthrax toxin undergoes a prepore to pore conformational change under acidic conditions. Previous approaches have generally used urea to avoid aggregation during the transition from PA_prepore_ to PA_pore_ (16–21). These approaches have limitations in that they do not account for the low pH electrostatic microenvironment in the pore lumen predicted to be important for LF-PA interactions (22) and they assume similar outcomes for chaotrope and acid induced unfolding. In order to overcome these limitations we have developed a novel assembly method for toxin immobilization, translocation, and nanodisc stabilization (called TITaNS, **Fig. 1B-G**). TITaNS was designed to mimic important low pH physiological states during the anthrax intoxication mechanism. This approach allows for endosomal pH pore formation and imaging of individual complexes in a lipid bilayer in the biologically relevant low pH environment (16, 23). TITaNS can be used in combination with techniques other than cryoEM, including mass spectrometry, nuclear magnetic resonance, surface plasmon resonance, and biolayer interferometry. TITaNS also has the potential to be adapted to screen prospective pharmaceuticals that arrest or prevent endosomal membrane insertion (23).

Reversible immobilization is key to the TITaNS methodology, because it allows the stabilized complexes to be released into solution. We began with recombinantly purified, soluble forms of LF_N_ and PA_prepore_ that are mixed together in solution. The binary complex of LF_N_ bound to PA_prepore_ was then immobilized onto thiol sepharose beads by covalently coupling E126C LF_N_ to the bead surface (**Fig. 1B**). The immobilized LFN-PA_prepore_ complex was oriented such that when the bead slurry was washed in pH 5.5 buffer to transition PA from prepore to pore, the pore extended away from the bead surface (**Fig. 1C**). We predict this low pH environment initiates translocation of LF_N_ through PA_pore_ *in vitro* (**Fig. 1D**). We base this prediction on computational and experimental evidence of early translocation events induced by low pH. Specifically, molecular simulations of anthrax toxin early translocation events predict the events are strongly influenced by the protonation state of LF and are highly favorable at low pH (22). Also at low pH, the partial translocation of LF has been observed in planar lipid bilayers (9). After pore formation, the next step in TITaNS was the stabilization of LF_N_-PA_pore_ translocation complexes using nanodisc technology (24–26). Pre-nanodisc micelles were added to the bead slurry and associated with the transmembrane portion of PA_pore_ (**Fig. 1E**). To promote lipid bilayer formation, we dialyzed away excess detergent (**Fig. 1F**). The soluble LFN-PA_pore_-nanodisc complexes were then eluted off the thiol sepharose beads using the reducing agent dithiothreitol. Eluted complexes were transferred to the cryoEM grid and the pH was dropped to pH 5.5 to capture the complex at low pH prior to blotting and plunge freezing (**Fig. 1G**). To prevent aggregation and migration of the nanodiscs to the air-water interface, we plunge froze the grids within 30 seconds of sample application. Using our TITaNS methodology, we obtained a 3.3 Å reconstruction of LF_N_ translocating through PA_pore_ (**Fig. 2A-D)**.

**Fig. 2.**
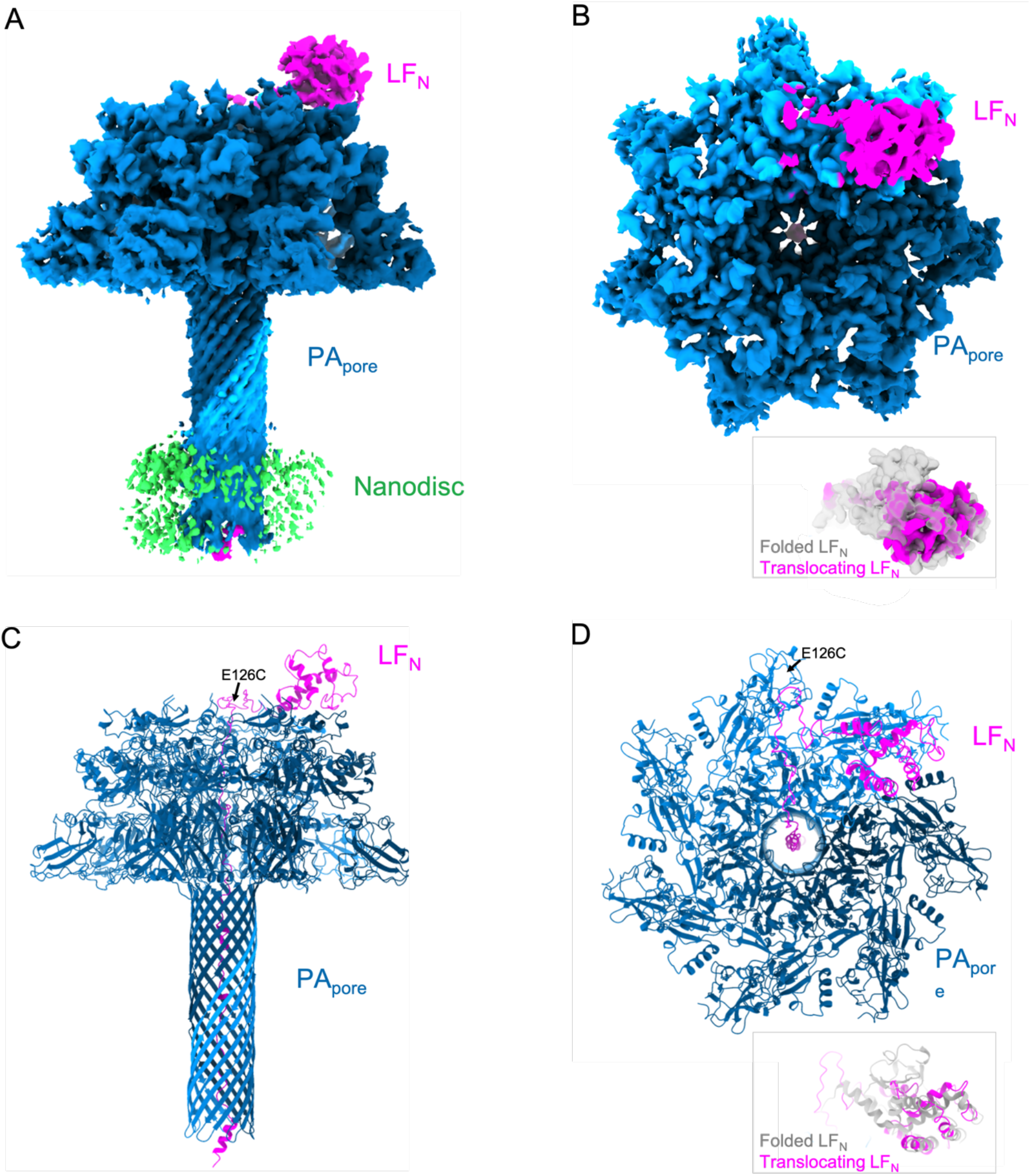
Overview of the anthrax toxin translocation complex. CryoEM density map (A) side and (B) top view of LF_N_ translocating through PA_pore_. Inset: folded LF_N_ model density (PDB 6PSN) compared to translocating LFN density. Molecular model (C) side and (D) top view of LFN translocating through PA_pore_. Inset: folded LF_N_ model (PDB 6PSN) compared to translocating LF_N_ model.

### Comparison of LF_N_ before and during Translocation

Prior to translocation, LF_N_ is bound to the cap of PA at the interface of two PA protomers with helix α1 bound to the α clamp (11, 17). Our reconstruction shows a loss of LF_N_ density in the canonical binding site above the PA_pore_ (**Fig. 2A,B**). Lethal factor translocates through PA_pore_ from the N-terminus to the C-terminus. Notably, our reconstruction does not show density for residues 50-135 in the canonical folded LF_N_ position (**Fig. 2B,D)** (11, 16). However, density for residues 136-250 are apparent. Our model of unfolding, translocating LFN is consistent with a molten globular intermediate state at low pH (27). In our translocating LF_N_ complex, immobilized residue E126C has not yet translocated suggesting translocation occurred while the residue was coupled to the bead surface at pH 5.5, stalling the complex at this position (**Fig. 2C,D**).

### Molecular Interactions of Translocation Complex

In addition to a loss of LF_N_ density in the canonical binding site above the PA_pore_, our reconstruction shows added density inside the pore lumen indicating a translocating complex **(Fig. 3)**. Density near the top of the PA_pore_ funnel was in proximity to several hydrophobic residues of PA_pore_ (**Fig. 3B**). Some of these residues, including Phe202, Phe236 and Phe464 are in the α clamp and have previously been proposed to aid in unfolding LF and stabilizing unfolded intermediates as they transition into the pore (11). Additional hydrophobic residues further down the pore lumen, such as Trp226 and Tyr456 may also help to stabilize the unfolded peptide. Our results are consistent with these hydrophobic residues facilitating translocation of LF_N_ into the pore and toward the Φ clamp in an unfolded state. We observed added asymmetric density in and around the Φ clamp (**Fig. 3C**) that was not visible when we compared it to the previously published apo PA_pore_ cryoEM structure (13). Specifically, there is density in the center of the Φ clamp (**Fig. 3C**). We attribute this density to unfolded LF_N_ interacting with the Phe ring of the PA_pore_ Φ clamp loops as LF_N_ is translocating through the pore. In addition the density for each of the PA_pore_ Phe427 residues was smeared in plane with the benzyl ring suggesting rotameric states moving up and down (**Fig. 3D)**.

**Figure 3.**
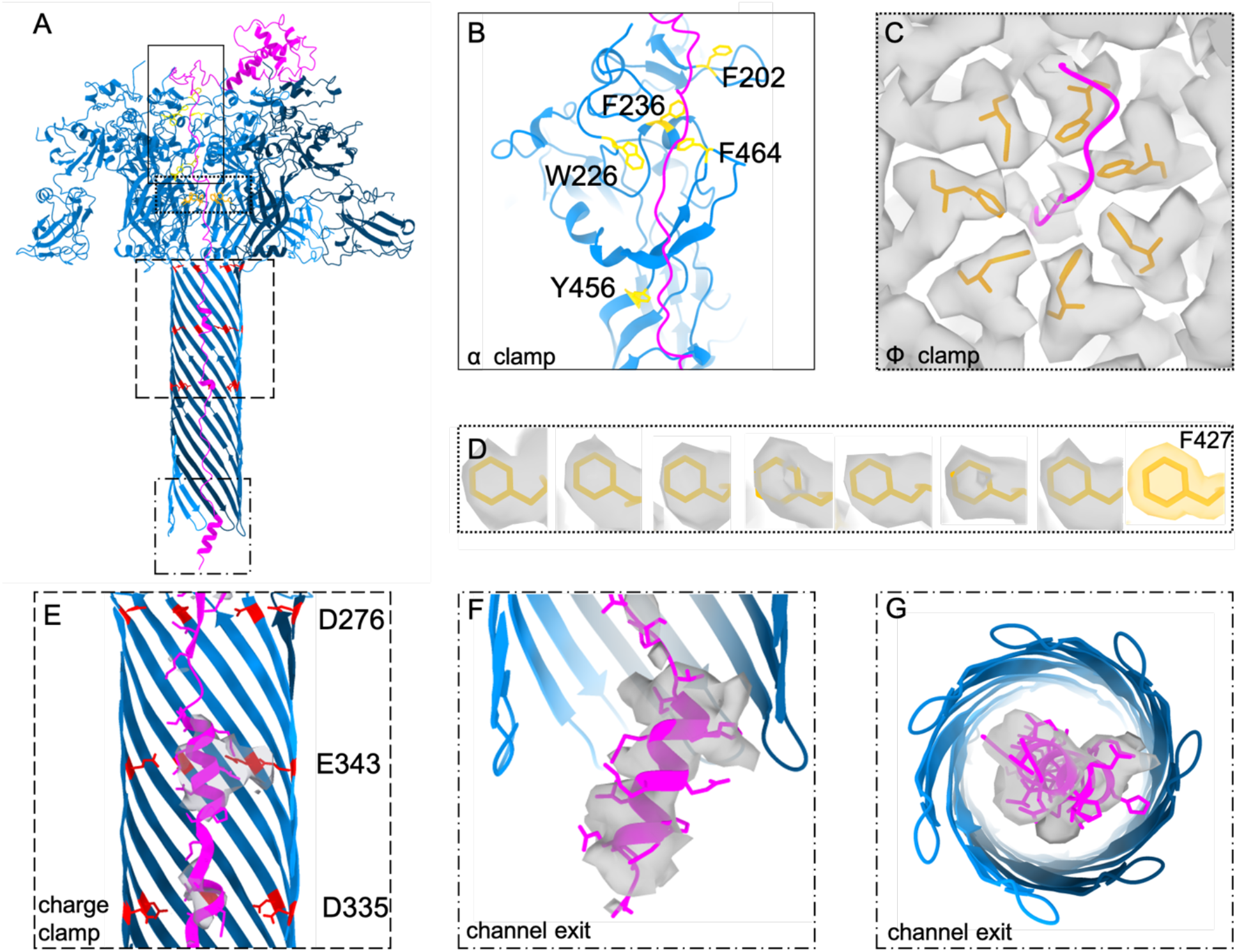
Unfolding and refolding of LF_N_ at key clamp sites during translocation through PA_pore_. (A) Model of PA_pore_ (blue) translocating LF_N_ (magenta) with α clamp, Φ clamp, and charge clamp residues shown in yellow, orange, and red respectively. (B) Hydrophobic residues predicted to facilitate unfolding in early translocation from α clamp to Φ clamp. (C) Φ clamp ring with LF_N_ density. (D) Individual F427 residues (orange) with associated cryo-EM density (grey) for each subunit compared to modelled density (orange). (E) LFN model (magenta) and density (grey) in β barrel charge clamp (F) PA_pore_ channel exit into the cytosol with LF_N_ α1 helical model (magenta) and density (grey). (G) 90° rotation of F.

Translocating LF_N_ density was also observed in the β barrel of the PA_pore_. Focused refinement of the β barrel interior revealed density consistent with α helices along with portions of unfolded peptide (**Fig. 3E-G)**. Notably, density located at the PA_pore_ charge clamp is consistent with an α helix and suggests the deprotonated state of LF favors helix formation within the pore. Canonical charge clamp residues Asp276, Glu343, and Asp335 are shown in **Fig. 3E** with the predicted LF_N_ density translocating through the center of the channel. Further down the pore we observe density for helix α1 emerging from the channel exit **(Fig. 3F,G)**. Our results provide evidence for initial refolding of LF secondary structure both inside and upon exit from the PA_pore_. These results are consistent with ribosome exit tunnel studies that, using optical tweezers and molecular dynamics simulations, showed excluded volume effects and electrostatic interactions contribute to substrate folding (28).

Next, we wanted to determine whether there were any differences between PA_pore_ protomers that interacted directly with the translocating LF_N_ compared to protomers that did not interact with translocating LF_N_. Importantly, symmetry operations were not used during single particle analysis. To compare the PA_pore_ protomers, we aligned the seven protomer chains that compose the PA_pore_ **(Fig. S1A)**. Comparison of the chains showed little difference in the backbone or side chain rotamers for the majority of the residues at the LF_N_-PA_pore_ interfaces as well as the protomer-protomer interfaces. This rigidity is likely necessary for the PA_pore_ to maintain a stable β barrel and perform its function under endosomal conditions. During this analysis, we also examined the conformation of the receptor binding domain of PA_pore_. The receptor binding domain is connected to the main body of PA_pore_ by a single loop and is responsible for anchoring the toxin to the host cell membrane prior to complex endocytosis and pore formation (29). Interestingly, the receptor binding domain did show different conformations for each protomer and indicates a degree of conformational flexibility **(Fig. S1A-C)**. This variability between receptor binding domains likely arose from the acidic conditions and lack of a receptor. Overlaying our PA_pore_ chains with the crystal structure of PA_prepore_ bound to its receptor, capillary morphogenesis protein 2 (CMG2), revealed several conformations not conducive to receptor binding **(Fig. S1C)**. Specifically, PA E194 was not in position to form the metal ion-dependent adhesion site (MIDAS) motif in the receptor binding pocket (30). This indicates that without a receptor bound, the receptor binding domain loops adopt multiple states.

## Discussion

The anthrax toxin PA_pore_ unfolds, translocates, and refolds LF, it’s enzymatic subtrate. Three clamp sites aid in peptide translocation: the α clamp, the Φ clamp, and the charge clamp. We report here, cryoEM density consistent with nascent polypeptide chain translocating the length of PA_pore_. In our model, LF_N_ can be seen unfolding prior to the α clamp, passing through the dynamic Φ clamp, and refolding in the β barrel channel. This model is consistent with LF needing to completely unfold in order to translocate. However, if the entire 90 kDa enzyme were to unfold at once, deleterious folded intermediates or aggregates would likely block the PA_pore_ translocon, especially when multiple LF are bound to PA. Therefore, in order to efficiently translocate and refold, LF unfolds from the N to C terminus (31). While the low pH of the endosome destabilizes the enzyme, it does not completely unfold into its primary sequence (27, 32). Our results are consistent with molten globular translocation intermediates of LF being destabilized in the acidic environment of the endosome (27) with the α clamp then able to apply additional unfolding force on the protein and funnel LF towards the Φ clamp (2). Translocation requires step-wise unfolding and stabilization of the unfolded intermediates to prevent aggregation. When LF binds to PA_pore_, helix α1 of LF moves away from the main body of LF and binds to the α clamp of PA_pore_ (11). From here, LF has multiple paths it could take through the PA_pore_ funnel, gated by the Φ clamp. Our results suggest a favorable path from the α to Φ clamp that involves a series of hydrophobic residues that are amenable to unfolded translocation intermediates and likely serve as checkpoints to verify the unfolded state of LF prior to the Φ clamp (**Fig. 3A-B)**.

The Φ clamp plays a crucial roll in translocation by acting as a hydrophobic seal between the endosome and cytosol (9). Multiple Φ clamp states have been hypothesized at pH 5.5 (33). Our analysis did not reveal multiple distinct states. However, the smeared density in our structure is indicative of a dynamic clamp. Our density does not imply dilation of the clamp (33), so much as a up and down motion along the pore axis. This motion could be conserted or individual F427 residues moving to accommodate various translocating side chains. The compressive and tensile forces generated by the unfolding LF in the PA_pore_ funnel above and the refolding LF in the β barrel channel below may also contribute to this movement. However, too much flexibility or dialation would cause the seal at the Φ clamp to be lost. Therefore, this dynamic motion must still maintain the pH gradient between the endosome and cytosol, while accomidating any side chain, ensuring efficient translocation.

Helix formation inside the PA_pore_ β barrel has been hypothesized but, to our knowledge, never observed (13, 27). We report here, evidence of concomitant α helix formation and translocation inside the β barrel of the PA_pore_ (**Fig. 3E,F,G)**. We hypothesized that, along with changing the charge state of the peptide substrate, the charge clamp allows for a local folding environment within the PA_pore_. Helical portions of LF have previously been shown to dock into the α clamp, with the periodicity of these helices aiding in efficient unfolding of LF (34). We predict this periodicity is also important for hypothesized refolding of LF, beginning at the charge clamp. Our hypothesis is consistent with other anthrax toxin subtrates, such as LF_N_ fused to the catalytic chain of diphtheria toxin (LF_N_-DTA), which did not evolve to fold in the PA_pore_ channel. Interestingly, these non-native substrates require chaperones for enzymatic activity (35) indicating the DTA portion of these proteins do not form helices in the PA channel at optimal intervals. Our model is also reminesant of the ribosome, where helix folding in the exit tunnel aids in co-translational folding of native proteins (36). We predict helix folding in the PA_pore_ β barrel aids in co-translocational folding by temporally altering LF emersion from the tunnel allowing regions to fold into tertiary structures.

A proposed unfolding-refolding translocation model is shown in **Fig. 4A** starting with LF_N_ bound to the funnel rim of PA_pore_. LF_N_ is unfolded and funnelled towards the Φ clamp, aided by hydrophobic residues along the funnel slope. LF_N_ acidic residues are protonated in the acidic environment of the funnel. Completely unfolded LF_N_ then passes the Φ clamp. This ring of F427 residues remains restrictive enough to maintain a seal while accomdating translocation. As the channel widens in the charge clamp, acidic residues are deprotonated. Folding of α helical portions places mechanical force on the translocating peptide, contributing to efficient translocation, and overcoming local energy minimum that could otherwise stall the complex. The newly formed, secondary structure favors unidirectional translocation by discouraging retrograde transfer through the narrow Φ clamp resulting in natively folded LF in the host cell cytosol.

**Fig. 4.**
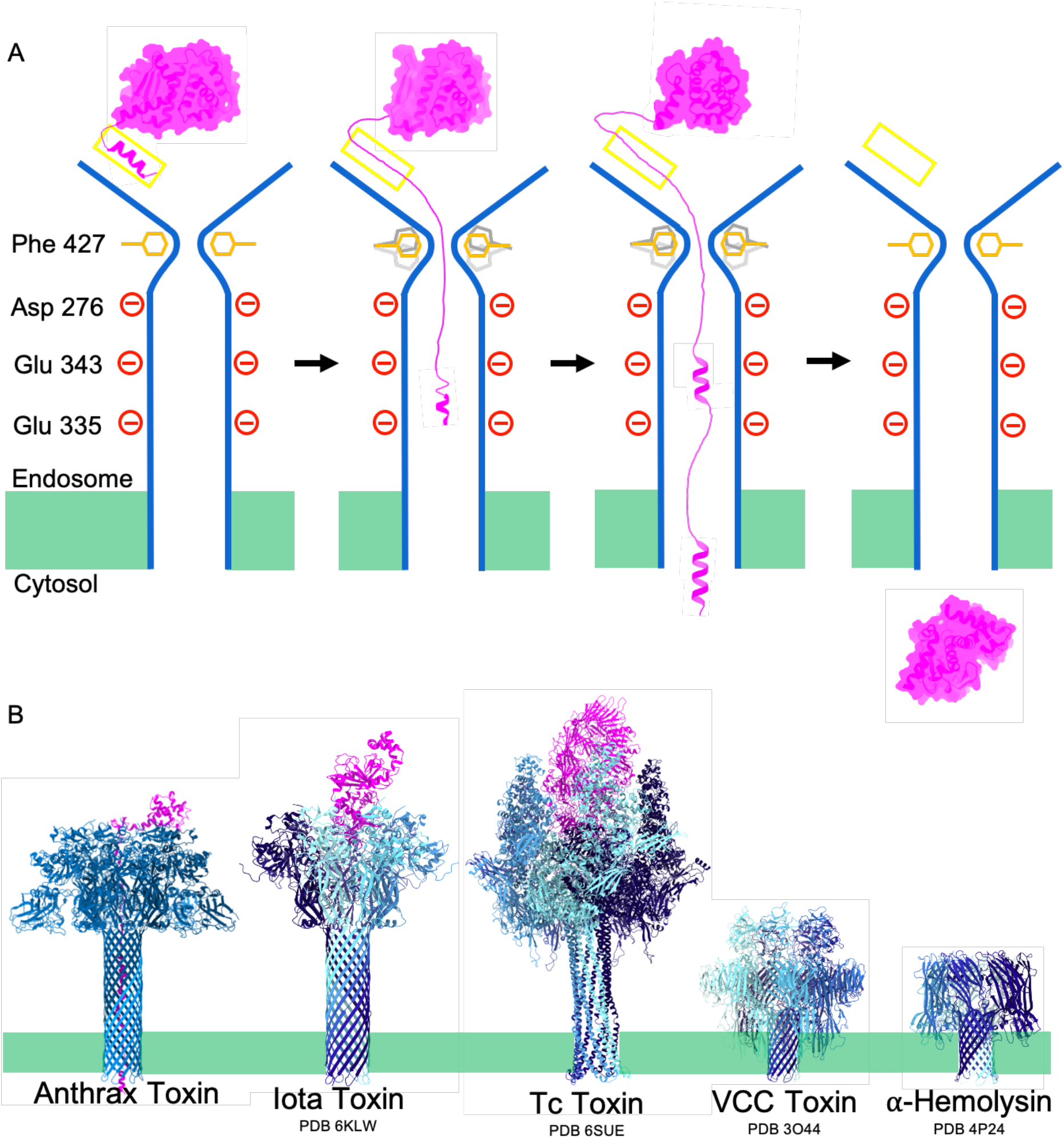
Proposed AB toxin extended pore refolding mechanism. (A) Unfolding-refolding translocation model of anthrax toxin. LF_N_ (magenta) translocates through PA_pore_ (blue) passing the α clamp (yellow), the Φ clamp (orange and grey), and the charge clamp (red). Helical portions of LF_N_ begin to refold in the channel ensuring proper tertiary refolding of LF_N_. (B) Comparison of toxin pore length between toxins that translocate proteins vs toxins that disrupt ion gradients. Membrane bilayer represented in green.

Our results have implications for other toxins. PA Phe427 is equivalent to Phe454 of *Clostridium perfringens* iota toxin (37), Phe428 of *Clostridium botulinum* C2II binary toxin (38), and Trp318 of *Vibrio cholerae* cytolysin (39). We hypothesize a dynamic hydrophobic seal model is a common mechanism, applicable to these other toxins. Initial refolding in the pore channel is also likely not unique to the anthrax toxin. Indeed, other translocons, such as the iota toxin and toxin complex (Tc) toxin (**Fig. 4B**), have pores that extend well passed the membrane bilayer (40, 41). We predict these pore forming toxins have evolved extended pores to faciliate substrate refolding inside the translocon for effective intoxication. Not all pore forming toxins translocate proteins. Some, like *Vibrio cholerae* cytolysin (VCC) and *Staphylococcus aureus* α-hemolysin, form pores to distrupt ion concentrations (39, 42). The pore length of these toxins is noticeably shorter (**Fig. 4B**).

## Materials and Methods

### Protein Expression and Purification

Proteins were purified as previously described (16). Briefly, His_6_-SUMO-LF_N_ E126C was expressed in BL21 cells, purified using anion exchange, and cleaved by small ubiquitin-related modifier protease (43). Recombinant wild-type PA_83_ was expressed in the periplasm of *Escherichia coli* BL21 (DE3) and purified by ammonium precipitation and anion exchange chromatography (8). After trypsin activation (43), PA_63_ heptameric prepores were formed using anion exchange and size exclusion chromatography. Membrane scaffold protein 1D1 (MSP1D1) was expressed from the pMSP1D1 plasmid (AddGene) with an N-terminal His-tag and was purified by affinity chromatography (26).

### LF_N_-PA-Nanodisc Complex Formation for CryoEM with TITaNS

E126C LF_N_ and PA_prepore_ were incubated in solution and then immobilized by coupling E126C LF_N_ to activated thiol sepharose 4B beads (GE Healthcare Bio-Sciences, Pittsburgh, PA, USA) in Assembly Buffer (50 mM Tris, 50 mM NaCl, 10mM CaCl_2_ pH 7.5) at 4 °C for 12 hr. Beads were washed three times with Assembly Buffer to remove any unbound PA_prepore_. The immobilized LFN-PA_prepore_ complexes were then incubated in low pH buffer (10 mM acetate, 50 mM Tris, 50 mM NaCl, 10mM CaCl_2_ pH 5.5) to transition the PA_prepore_ to PA_pore_ and initiate translocation of LF_N_. The beads were then washed in Assembly Buffer at neutral pH three times. Next, pre-nanodisc micelles (2.5 μM MSP1D1, 97.5 μM 1-palmitoyl-2-oleoyl-sn-glycero-3-phosphocholine (POPC) (Avanti, Alabaster, AL, USA), 65 (POPG) in 25 mM Na-cholate (Sigma-Aldrich, St. Louis, MO, USA), 50 mM Tris, and 50 mM NaCl) were added and bound to the aggregation-prone hydrophobic transmembrane β-barrel of PA_pore_. The micelles were collapsed into nanodiscs by removing Na-cholate using dialysis with Bio-Beads (BIO RAD, Hercules, CA, USA). Stabilized complexes were released from the thiol sepharose beads by reducing the E126C LFN-bead disulfide bond using 50 mM dithiothreitol (DTT) (Goldbio, St. Louis, MO, USA) in Assembly Buffer. Assembled complexes were initially confirmed using negative-stain TEM. Complexes were stored at −80C prior to cryoEM grid preparation. Complex formation has also previously been confirmed using mass spectrometry and biolayer interferometry (44).

### Grid Preparation for CryoEM

Complexes stored at −80C were thawed on ice. A glow discharged Quantifoil R1.2/1.3 300M Cu holey carbon grid was placed inside the FEI Vitrobot Mark IV humidity chamber at 100% humidity. Then, 2ul of thawed sample was applied to the grid followed by 0.5uL of 1M acetate pH 5.5. The grids were then blotted and plunge frozen in liquid ethane. Frozen grids were stored in liquid nitrogen prior to use.

### CryoEM Data Collection and Image Processing

CryoEM grids were loaded into a FEI Titan Krios electron microscope operated at 300 kV for automated image acquisition with serialEM (45). cryoEM micrographs were recorded as movies on a Gatan K2 Summit direct electron detection camera using the electron counting mode in super resolution mode at ×130K nominal magnification, a pixel size of 0.535 Å per pixel, and defocus ranging between −1 and −3 μm. Total dose was 50.76 e-/ Å ^2^. Total exposure time was 9s and fractionated into 45 frames with 200 ms exposure time for each frame. In total, 6,515 micrographs were taken in a continuous session. Frames in each movie were aligned and averaged for correction of beam-induced drift using MotionCor2 and cryoSPARC patch motion correction to generate a micrograph (46, 47). Micrographs generated by averaging all frames of each movie were used for defocus determination and particle picking. Micrographs obtained by averaging frames 2-36 (corresponding to ~40 e/Å^2^) were used for two- and three-dimensional image classifications. The best 4,488 micrographs were selected for the following in-depth data processing.

### Single Particle Analysis and Density Modification

Single particle analysis was performed using cryoSPARC v2.15 (46) **(Fig. S2)**. A random subset of micrographs was selected for blob particle picking. These particles were subjected to 2D classification in order to obtain a set of five particle templates. Using these templates, 2,076,581 particles were selected from 4,488 micrographs. After multiple rounds of 2D classification, the remaining 671,090 ‘good’ particles were used to create an *ab initio* model. Heterogenous classification with four classes was then performed and the class with full length β barrel and nanodisc was selected. To select particles with LF_N_, 3D variability analysis was performed with three orthogonal principle modes (i.e. eigenvectors of 3D covariance) and a mask of LF_N_ bound in each of the seven possible binding sites filtered to 30 Å and gaussian blurred. 122,651 particles from three resulting clusters with potential LF_N_ density were selected for further processing. A non-uniform refinement on the per particle motion and CTF corrected particles was performed resulting in a 3.3 Å cryoEM density map. Resolution was determined using gold standard Fourier shell correlation with a cut off of 0.143. Next, local refinements and density modifications were performed (**Fig. S3**). To further characterize bound LF_N_, local refinement of the cap of the PA_pore_ was performed using a mask of LF_N_ (PDB 3KWV) low pass filtered to 30 Å. For the β barrel interior, a cylindrical mask was used. Phenix density modification was perfomed on the non-uniform refinement and local refinement half maps to further improve the density for PA_pore_ and LF_N_, respectively (48). Phenix combined focused maps was then used to create a composite map (**Fig. S3**) (49). The map showed density surrounding the transmembrane region of the beta barrel that we interpret as nanodisc density. As has been reported previously (13), there was also disordered density surrounding the outside of the middle of the β barrel. This additional density was masked out of the final map using a 30 Å mask of LFN-PA_pore_ inserted into a nanodisc.

### Model Building and Refinement

An initial model using PDB 6PSN was docked into the cryoEM map using Chimera map to model (50). The LF_N_ coarse model was adjusted manually using Coot (51) to fit the density starting at the C-terminus. Model α helical assignments were based on helicity in original model, cryoEM density diameter, and consistency with previously published helical density. The PA_pore_ coarse model was refined using PHENIX real space refine (49). Individual atomic model side chains were manually adjusted to fit the density map using Coot (51). This process was repeated iteratively until an optimal model was obtained. Ramachandran plots and MolProbity (52) were used to assess model quality. **Supplementary Table 1** is a summary of cryoEM data collection and processing as well as model building and validation.

## Acknowledgements

This work is dedicated to the memory of our friend, mentor, and colleague Dr. Mark T. Fisher. This work was supported by National Institutes of Health (R35-GM128562 and R03-AI142361 to B.D.F.), and by University of Kansas Madison and Lila Self Graduate Fellowship to A.J.M. The authors acknowledge the use of instruments at the Electron Imaging Center for NanoMachines supported by NIH (1S10RR23057 and 1S10OD018111), NSF (DBI-1338135) and CNSI at UCLA.

---

**Fig. S1.**
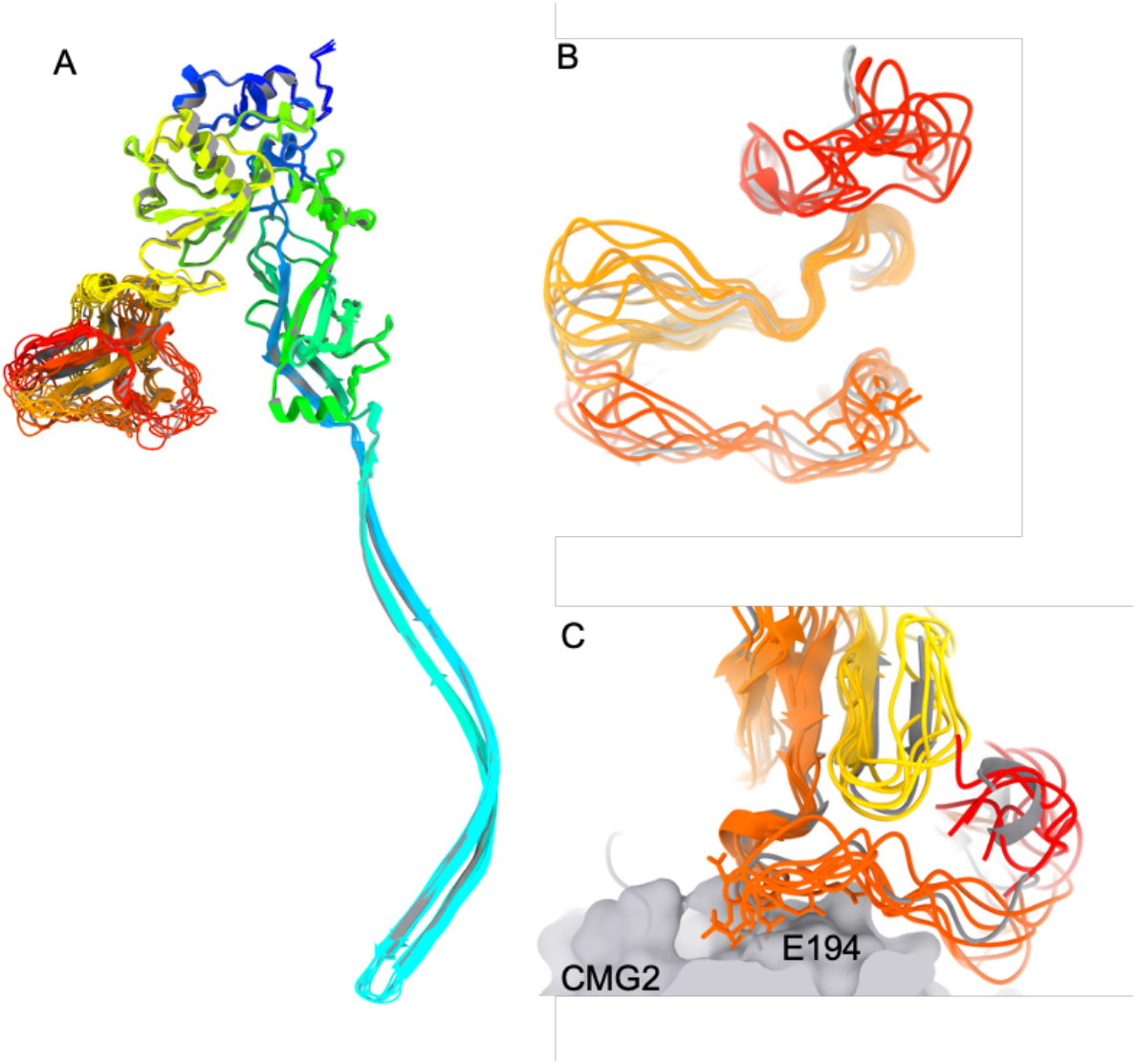
Comparison of PA_pore_ protomers. (A) PA_pore_ protomer chains are shown in rainbow N to C termini for chains A-G. The receptor binding domain is shown in yellow, red, and orange. (B) Bottom up view of receptor binding domain with PA_prepore_ receptor binding domain shown in grey (PDB 1T6B). (C) Comparison of PA_pore_ protomers to PA_prepore_ bound to receptor CMG2 (grey).

**Fig. S2.**
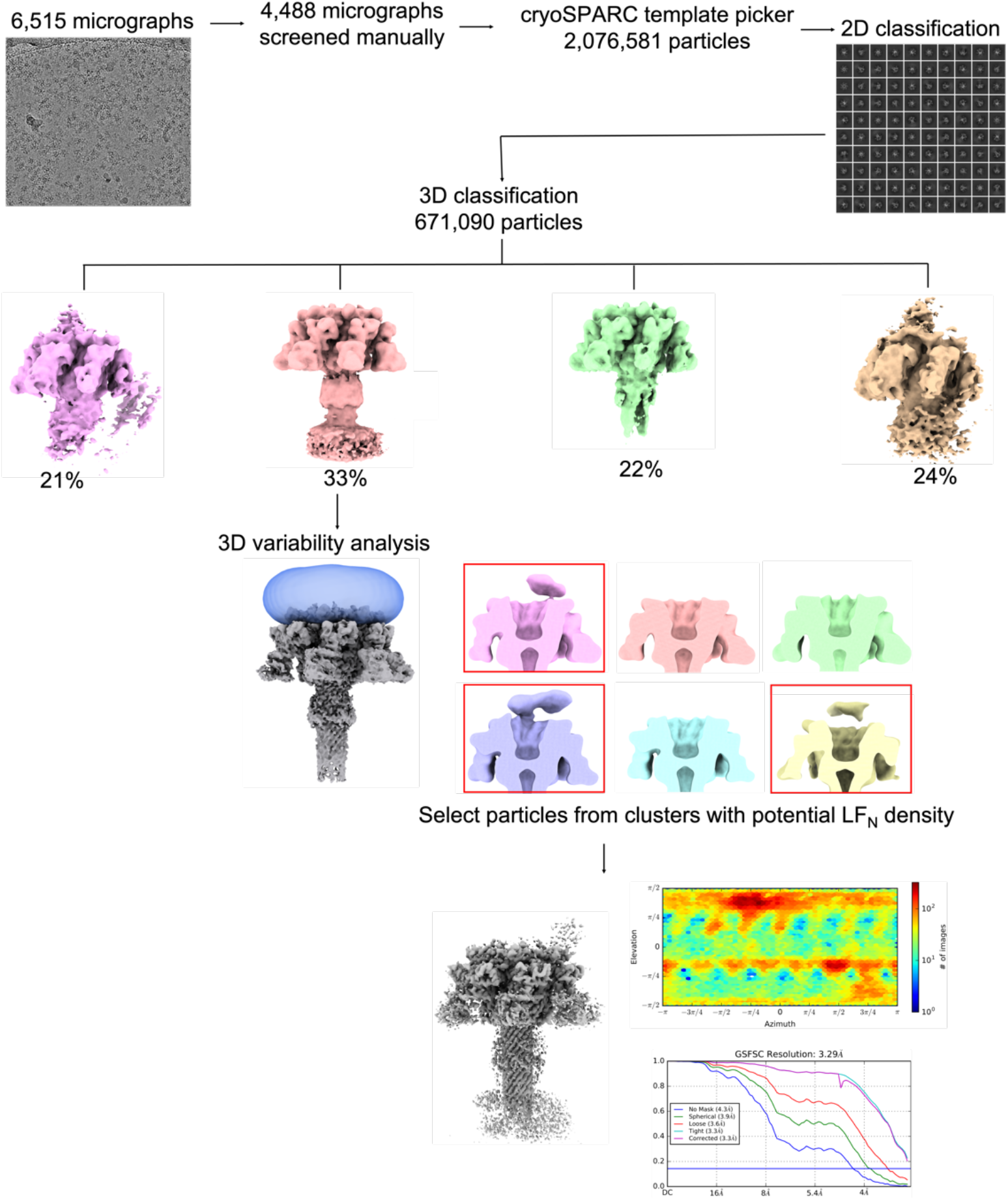
Single particle analysis of anthrax toxin translocating complexes. After micrograph curation and 2D classification, ‘good’ particles were subjected to 3D classification (four classes) followed by 3D variability analysis on particles that contained intact β barrel. 3D variability analysis focused on potential LF_N_ binding sites (blue mask) above PA_pore_ (grey). Results of 3D variability analysis are shown in pastels filtered to 20 Å (pink, orange, green, purple, blue, yellow) with potential LF_N_ density containing maps boxed in red. 3.3 Å Non-uniform refinement of PA_pore_ with intact β barrel and potential LF_N_ density is shown in grey (FSC cutoff 0.143). Euler angle distribution of particles shows diverse particle orientations.

**Fig. S3.**
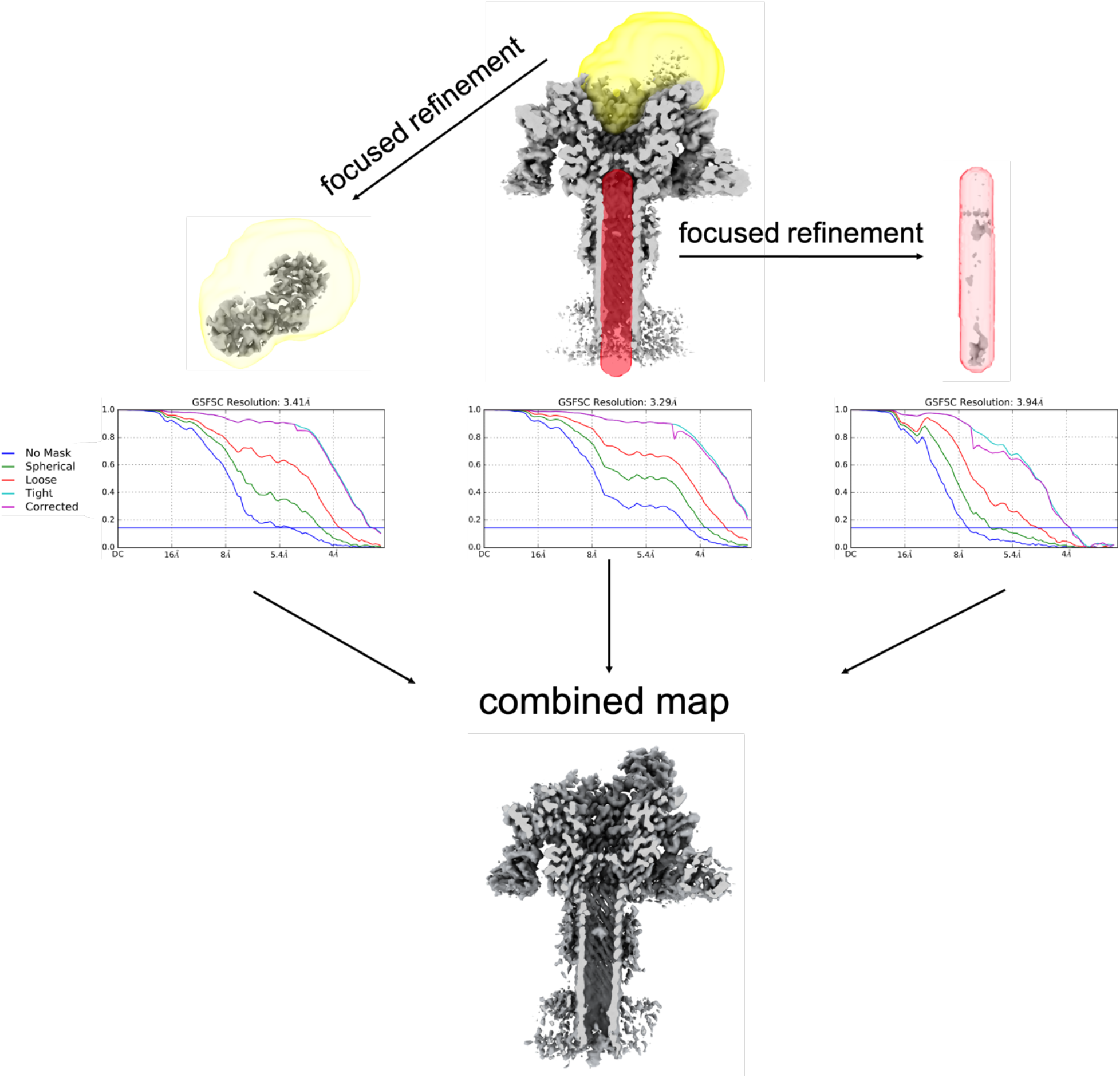
Density modification of cryoEM map. Focused refinement and density modification of non-uniform refined cryoEM map were perfomed in cryoSPARC and Phenix, respectively. Density for LF_N_ above PA_pore_, PA_pore_, and β barrel interior were combined using Phenix combine focus maps. Resolution was determined for each density prior to combining using FSC cutoff of 0.143.

**Table S1.** cryoEM Data Collection and Processing Statistics.

|  |  |
| --- | --- |
| **Data collection and processing** |  |
| Magnification | 130,000X |
| Voltage (kV) | 300 |
| Electron exposure ($e^-/\text{\AA}^2$) | 1.79 per frame |
| Defocus range ($\mu\text{m}$) | 1-3$\mu\text{m}$ |
| Pixel size ($\text{\AA}$) | 0.535 |
| Symmetry imposed | C1 |
| Initial particle images (no.) | 2076581 |
| Final particle images (no.) | 122651 |
| Map resolution ($\text{\AA}$) | 3.3 |
| FSC threshold | 0.143 |
| **Refinement** |  |
| Initial model used (PDB ID) | 6PSN |
| Model resolution ($\text{\AA}$) | 3.3 |
| FSC threshold | 0.143 |
| **Model composition** |  |
| Nonhydrogen atoms | 32826 |
| Protein residues | 4154 |
| Ligands | 14 |
| **B factors ($\text{\AA}^2$)** |  |
| Protein | 395 |
| Ligand | 202 |
| **r.m.s. deviations** |  |
| Bond Length ($\text{\AA}$) ($\# > 4\sigma$) | 0.009 (2) |
| Bond Angles ($^\circ$) ($\# > 4\sigma$) | 1.311 (37) |
| **Validation** |  |
| MolProbity score | 2.75 |
| Clashscore | 26 |
| Poor rotamers (%) | 4.1 |
| **Ramachandran plot** |  |
| Favored (%) | 94.25 |
| Allowed (%) | 5.32 |
| Disallowed (%) | 0.43 |

